# ZFC3H1 and U1-70K promote the nuclear retention of mRNAs with 5’ splice site motif within nuclear speckles

**DOI:** 10.1101/2021.06.08.447610

**Authors:** Eliza S. Lee, Harrison W. Smith, Eric J. Wolf, Aysegul Guvenek, Andrew Emili, Bin Tian, Alexander F. Palazzo

## Abstract

Quality control of mRNA represents an important regulatory mechanism for gene expression in eukaryotes. One component of this quality control is the nuclear retention and decay of misprocessed RNAs. Previously, we demonstrated that mature mRNAs containing a 5’ splice site (5’SS) motif, which is typically found in misprocessed RNAs such as intronic polyadenylated (IPA) transcripts, are nuclear retained and degraded. Here we demonstrate that these transcripts require the zinc finger protein ZFC3H1 for their decay and nuclear retention into nuclear speckles. Furthermore, we find that U1-70K, a component of the U1 snRNP spliceosomal complex, is also required for their nuclear retention and likely functions in the same pathway as ZFC3H1. Finally, we show that the disassembly of nuclear speckles impairs the nuclear retention of mRNAs with 5’SS motifs. Together, our results suggest a model where mRNAs with 5’SS motifs are recognized by U1 snRNP, which then acts with ZFC3H1 to both promote their decay and prevent nuclear export of these mRNAs by sequestering them in nuclear speckles. Our results highlight a splicing independent role of U1 snRNP and indicate that it works in conjunction with ZFC3H1 in preventing the nuclear export of misprocessed mRNAs.

## Introduction

Eukaryotic cells are divided into two compartments, the nucleus and cytoplasm. In the nucleus, DNA is transcribed into pre-mRNAs, which undergo a number of processing events to generate mature mRNAs. These mRNA are then exported to the cytoplasm where they are translated into proteins. Importantly, a subset of all transcripts are misprocessed (Pickrell et al. 2010; Skandalis 2016) and if transported to the cytoplasm, they would be translated into truncated proteins that are often toxic (Veitia 2007). To deal with these misprocessed transcripts, eukaryotic cells have evolved quality control mechanisms that prevent these mRNAs from being exported and instead target these RNAs for degradation (Lee et al. 2015). For example, transcripts that contain retained introns are retained in the nucleus and targeted for decay (Palazzo and Lee 2018). Such RNAs still contain intron-associated *cis*-elements, which are normally removed during splicing. These include the elements that specify the exon-intron boundaries (the 5’ splice site (5’SS) and the 3’ splice site (3’SS) motifs) and the branchpoint. Other misprocessed transcripts may be generated when a cryptic 3’ cleavage signal within an intron is used. These Intronic <u>P</u>oly<u>a</u>denylation (IPA) transcripts also include an unutilized 5’SS motif present near the 3’ end of the RNA (Tian et al. 2007). Interestingly, it has been proposed that many lncRNAs are also nuclear retained due to the fact that they have poorly spliced introns (Zuckerman and Ulitsky 2019). Moreover, other studies have found that many lncRNAs are retained by the U1 snRNP (Azam et al. 2019; Lubelsky et al. 2021; Yin et al. 2020), the component of the spliceosome that recognizes 5’SS motifs which are likely found in lncRNAs with retained introns.

Previously, we showed that RNA s that contain a consensus 5’SS motif near their 3’ end are retained in the nucleus and degraded (Lee et al. 2015). Our findings are consistent with other lines of experiments that have demonstrated that the presence of a 5’SS motif in a mature mRNA inhibits protein expression (Abad et al. 2008; Blázquez and Fortes 2013; Boelens et al. 1993; Fortes et al. 2003; Goraczniak et al. 2009; Guan et al. 2007; Gunderson et al. 1998; Vagner et al. 2000). It remains unclear how the 5’SS motif triggers nuclear retention. One possibility is that the U1 snRNP itself may play a role. This is in line with the finding that tethering U1 components to a reporter promotes nuclear retention (Takemura et al. 2011). Interestingly, U1 snRNP is present at very high levels, about one order of magnitude higher that other snRNPs (Baserga and Steitz 1993), supporting the idea that it has additional roles beyond splicing. The U1 snRNP components U1A and U1-70K have been shown to be required for the ability of the 5’SS motif to inhibit protein expression by repressing 3’ cleavage and efficient polyadenylation (Boelens et al. 1993; Gunderson et al. 1998; Vagner et al. 2000). The U1 snRNP is also required for the 5’SS motif to inhibit cleavage at cryptic 3’ cleavage signals in introns that produce IPA transcripts (Berg et al. 2012; Kaida et al. 2010). Other cryptic transcripts, especially upstream anti-sense transcripts from bi-directional promoters, often fail to recruit U1 and are rapidly cleaved and targeted for decay (Almada et al. 2013).

Beyond the U1 snRNP, it has been unclear what other factors may be involved. One likely candidate is the PAXT complex, which is composed of MTR4, ZFC3H1, PABPN1, and other accessory proteins (Meola et al. 2016; Ogami and Manley 2017; Ogami et al. 2017; Silla et al. 2020; Wu et al. 2020). Depletion of MTR4 or ZFC3H1 in Hela cells led to the accumulation of IPA transcripts in polysomes (Ogami and Manley 2017). Depletion of PAXT components also led to the upregulation of certain non-coding transcripts, such as a subset of enhancer RNAs (eRNAs) and <u>PRO</u>moter <u>UP</u>stream <u>T</u>ranscripts (PROMPTs) (Meola et al. 2016; Silla et al. 2020; Wu et al. 2020; Fan et al. 2017). MTR4 and ZFC3H1 may directly compete with the nuclear export adapter Aly (also known as AlyRef) for RNA binding (Fan et al. 2017; Silla et al. 2018), although whether this is related to any putative U1-mediated nuclear retention remains unclear.

Previously we noted that reporter RNAs that contained 5’SS motifs accumulated in nuclear speckles (Lee et al. 2015). These liquid-liquid phase separated nuclear compartments contain splicing factors, such as SR-proteins, nuclear export factors, and other mRNP factors such as components of the TREX and exon junction complexes (Akef et al. 2013; Daguenet et al. 2012; Dias et al. 2010; Dufu et al. 2010; Galganski et al. 2017; Schmidt et al. 2006; Spector and Lamond 2011). Although the majority of splicing occurs co-transcriptionally, certain post-transcriptional splicing is thought to occur in nuclear speckles (Girard et al. 2012; Mor et al. 2016). In support of the idea that post-transcriptional splicing occurs in nuclear speckles, it has been observed that certain reporter mRNAs expressed from plasmids are spliced in these structures (Dias et al. 2010).

How these mRNAs are targeted to speckles remains unclear. It has been observed that exon splicing enhancer elements, which recruit SR splicing factors and promote splicing, also help to target mRNAs to speckles (Wang et al. 2018). Since the 5’SS motif also promotes speckle localization, it is likely that U1 snRNP may also promote this relocalization. Indeed, there are significant amounts of U1 snRNP stored and/or localized in these structures (Huang and Spector 1992). The idea that U1 snRNP directs mRNAs to speckles is consistent with the observation that inhibition of either U2 or U4 snRNPs, which would be required to complete post-transcriptional splicing, caused the accumulation of unspliced mRNAs in nuclear speckles (Hett and West 2014; Kaida et al. 2007). Once splicing is complete, nuclear speckle-associated mRNAs may require the RNA helicase activity of the TREX-component UAP56 to exit speckles and promote nuclear export (Akef et al. 2013; Dias et al. 2010; Hondele et al. 2019).

Here we dissect the requirement for reporter RNAs that contain a 5’SS motif to be retained in the nucleus and the role of nuclear speckles in this process. We find that upon ZFC3H1-depletion, there is a global upregulation of IPA transcripts which accumulates in the cytoplasm. In addition, we observe that there is a defect in the nuclear export of mRNAs with long 3’UTRs that are generated from distal 3’ polyadenylation/cleavage sites. Using reporter mRNAs, we demonstrate that ZFC3H1, but not other PAXT components are required for the nuclear retention of mRNAs with intact 5’SS motifs in U2OS cells. We also find that U1-70K, a component of the U1 snRNP is also require for their nuclear retention and likely acts in the same pathway as ZFC3H1. We demonstrate that ZFC3H1 is required for mRNAs with 5’SS motifs to be retained in nuclear speckles and that nuclear speckle disassembly prevents this retention. Our data provides novel insight into how misprocessed mRNAs, and likely how certain functional non-coding RNAs (ncRNAs), are localized to the nucleus.

## Results

### Depletion of ZFC3H1 leads to the stabilization and cytoplasmic accumulation of Intronic Polyadenylation (IPA) transcripts

Previously, it had been shown that upon depletion of MTR4 or ZFC3H1 in HeLa cells, Intronic <u>P</u>oly<u>a</u>denylation (IPA) transcripts become stabilized and then accumulate in the cytoplasm (Ogami et al. 2017). This work looked at the overall distribution of mRNAs by next-generation sequencing in MTR4-depleted cells and only specific IPA transcripts by PCR in ZFC3H1-depleted cells. Interestingly, MTR4-depletion in HeLa cells also led to ZFC3H1 co-depletion (Ogami et al. 2017; Silla et al. 2018), likely by destabilizing the PAXT complex. Thus, it was not clear whether the effects observed on IPA transcripts were due to the direct depletion of MTR4, the indirect depletion of ZFC3H1, or a combination of the two.

Other work with ZFC3H1-depleted or knockout cells examined whole transcriptome sequencing with respect to the decay and nuclear retention of PROMPTs and eRNAs, but not IPA transcripts (Meola et al. 2016; Silla et al. 2020; Wu et al. 2020). Furthermore, to infer effects on nuclear export, sequencing must be carried out on cytoplasmic and nuclear fractions independently, which has yet to be performed on cells lacking or depleted of ZFC3H1.

To investigate how ZFC3H1 affects the distribution and levels of various mRNAs, we depleted this protein using lentiviral delivered shRNA (“ZFC3H1-2”), isolated total, nuclear and cytoplasmic fractions and purified poly(A)-selected RNAs for next generation sequencing (Figure 1A). The fractions had minimal cross contamination (Figure 1B) and the depletion was almost complete (Figure 1C). Note that ZFC3H1-depletion had no observable effects on MTR4 levels (Figure 2A).

**Figure 1.**
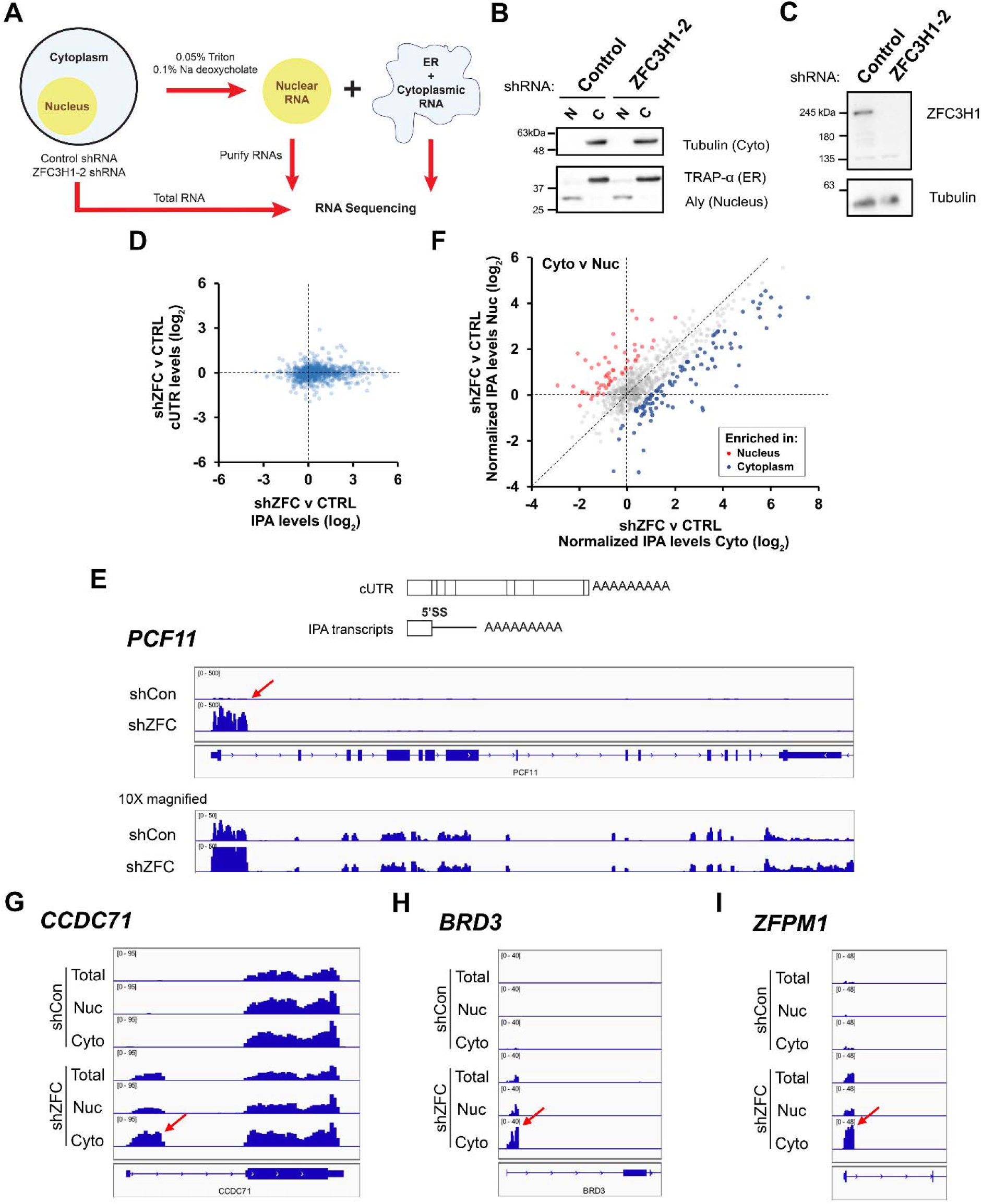
ZFC3H1-depletion leads to cytoplasmic accumulation of endogenous 5’SS motif containing mRNAs (or intronic polyadenylated transcripts) (A) Workflow for RNA Frac-Seq. ZFC3H1- or control-depleted U2OS cells were fractionated into nuclear and cytoplasmic/ER fractions, see Material and Methods for more details. RNA was purified from these fractions and from total cell lysates, and then analyzed by Illumina Sequencing. (B) Nuclear “N” and cytoplasmic “C” fractions were collected from ZFC3H1- or control-depleted U2OS cells, then separated by SDS-PAGE and analyzed by immunoblot for nuclear (Aly), ER (Trap-α) and cytoplasmic (Tubulin) protein markers. (C) Lysates collected from ZFC3H1- or control-depleted U2OS cells (96 hours post treatment with lentiviral delivered ZFC3H1-2 shRNA) were analyzed by immunoblot for ZFC3H1 and Tubulin. (D) Fold change in total levels of Intronic Polyadenylated (IPA) transcripts (ZFC3H1-depletion vs control depletion) (*x-axis*), plotted against the change in the total levels of fully processed mRNA (using cUTR reads, *y-axis*). Each dot corresponds to reads from one gene that is known to produce IPA transcripts (listed in Supplemental Table 1). Note that ZFC3H1-depletion leads to the upregulation of IPA transcripts, but not fully processed mRNAs. (E) Top, schematic of a fully processed mRNA and IPA transcript generated from the same gene. Note that the IPA transcript is generated from a 3’ cleavage/polyadenylation signal in the first intron and contains a 5’SS motif. Bottom, genome browser tracks of the *PCF11* gene in control-“shCon” or ZFC3H1-depleted “shZFC” cells at 500x and 50x resolution. Note the large peak for the IPA transcript (intronic cleavage/polyadenylation site is denoted with a red arrow), which is upregulated in ZFC3H1-depleted cells at 50x resolution. This transcript is also seen in control cells at higher magnification (500x). Also note the reads that correspond to the full-length transcript which are present in both control and ZFC3H1-depleted cells. (F) Similar to (D), except the fold change (ZFC3H1-depletion vs control depletion) in cytoplasmic levels of Intronic Polyadenylated (IPA) transcripts (*x-axis*) is compared to the fold change (ZFC3H1-depletion vs control depletion) in nuclear fraction (*y-axis*). Note that ZFC3H1-depletion leads to cytoplasmic accumulation of IPA transcripts (compare blue dots to red). To account for reads from fully processed mRNAs, the IPA transcript levels are normalized to the cUTR transcript levels of the same gene. (G-I) Genome browser tracks of three IPA transcript-producing genes, “*CCDC71*”, “*BRD3*” and “*ZFPM1*”. Note the accumulation of IPA transcripts in the cytoplasmic fractions. The intronic 3’ cleavage/polyadenylation sites are denoted by red arrows.

**Figure 2.**
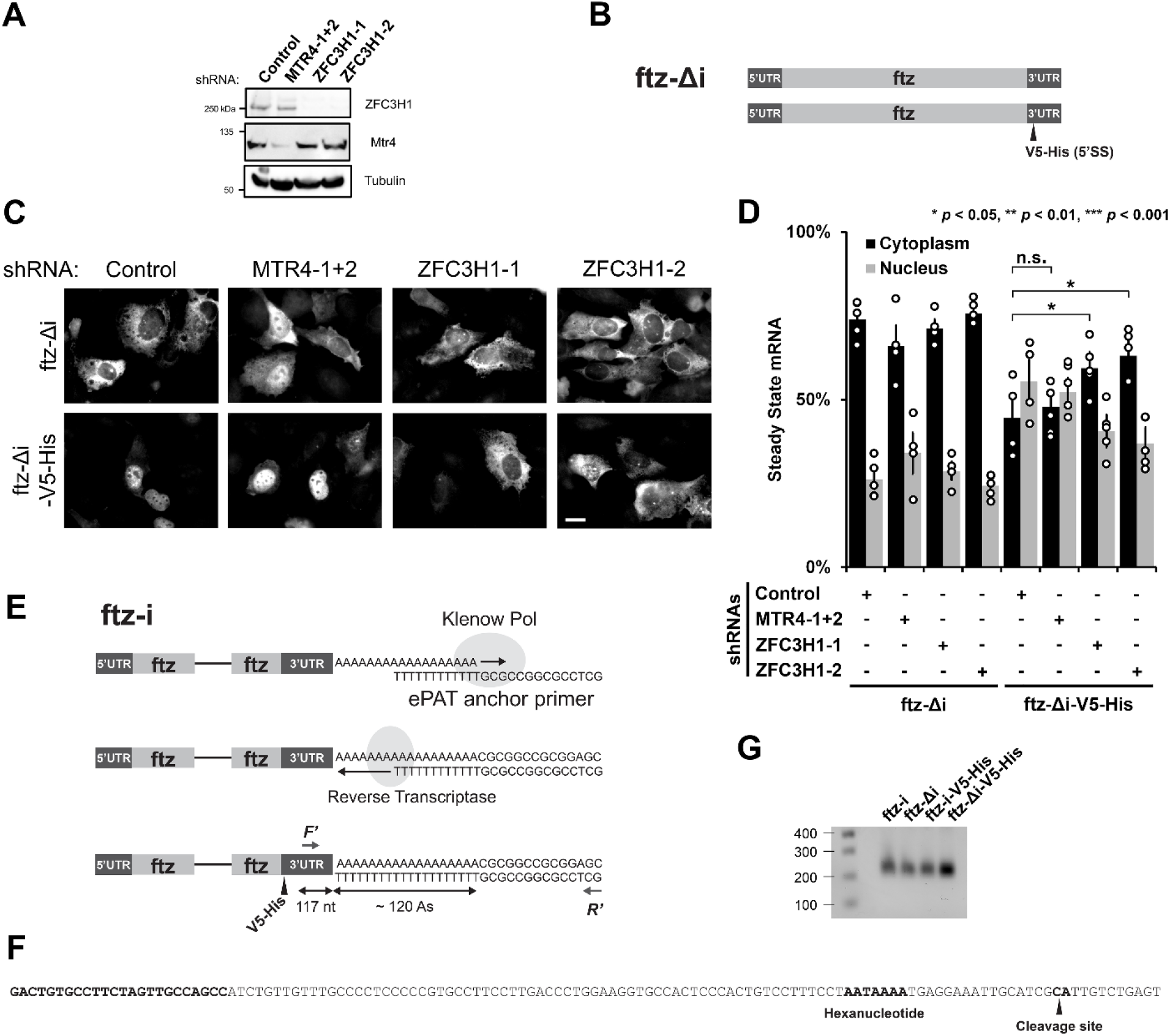
ZFC3H1 is required for the nuclear retention of 5’SS motif containing mRNAs. (A) U2OS cells were treated with different lentivirus shRNAs against ZFC3H1 (“ZFC3H1-1” and “ZFC3H1-2”), MTR4 (“MTR4-1+2”) or scrambled control. Lysates were collected after 96 hrs, separated by SDS-PAGE and immunoprobed for ZFC3H1, MTR4 or tubulin. Note that to effectively deplete MTR4, cells were treated with lentivirus containing both shRNA plasmids. (AB Schematic of the intronless (*Δi*) *ftz* reporter construct used in this study, with and without the *V5-His* element in the 3’ UTR. Note that the *V5-His* element contains a consensus 5’SS motif which promotes nuclear retention. (C-D) Control-, MTR4- and ZFC3H1-depleted cells were transfected with the intronless *ftz* reporter plasmid (+/- *V5-His*). 18-24 hours later the cells were fixed and the mRNA was visualized by FISH. Note that depletion of ZFC3H1, but not MTR4, caused cytoplasmic accumulation of the *ftz-Δi-V5-His*. Representative images are shown in (C), scale bar = 10 µm, and quantification is shown in (D). Each bar represents the average and standard error of at least three independent experiments, each experiment consisting of at least 30 to 60 cells. Student t-test was performed for Figure 2C, * *p* < 0.05, ** *p* < 0.01, *** *p* < 0.001. (E-G) ePAT assay was performed to examine 3’ end processing. (E) Schematic of the ePAT assay as described in Janicke et al 2013. Arrows represent the *ftz* specific F’ and universal R’ primers used to amplify the ePAT amplicon. The sequence of the ePAT amplicon before the cleavage site is shown in (F) and is 117 nucleotide long. (F) The sequence of the end of the 3’UTR is shown. Indicated in bold are the *ftz* specific F’ primer annealing site (used in the ePAT and 3’RACE experiments), the hexanucleotide motif, and the cleavage site (as determined by 3’RACE experiments on mRNAs derived from U2OS cells transfected with either *ftz-Δi* or *ftz-Δi-V5-His*). (G) PCR products from the ePAT assay were separated on a 1% agarose gel and stained with ethidium bromide. Lane 1: Molecular weight markers with sizes in bp indicated on the left, lanes 3-6: ePAT amplicons from U2OS cells that were transfected with plasmids containing the indicated versions of the *ftz* reporter (without (*Δi*) or with (*i*) an intron, without or with the *V5-His* element). Note that the amplicons generated from all 4 reactions are the same length (∼ 250 nucleotides). Since the amplified region in the 3’UTR is 117 bp long (see F), and the universal primer has a 14 nucleotide extension (see F), the poly(A)-tail is estimated to be approximately 120 nucleotides long.

In line with what had been seen in MTR4-depleted cells, ZFC3H1-depletion caused an upregulation of IPA transcripts with no detectable effect on fully processed mRNAs produced from the same genes (Figure 1D, Supplemental Table 1). When we looked at particular examples, such as *PCF11*, which has a well characterized IPA (Wang et al. 2019), we observed that it was upregulated more than 10 fold after ZFC3H1-depletion (Figure 1E, note that the 3’ end of the *PCF11* IPA transcript is indicated by the red arrow). The upregulation of IPA transcripts was also apparent for other genes (Supplemental Figure 1A-C). Although most IPAs occur through the action of cryptic 3’ polyadenylation/cleavage sites in the first intron, a minority are generated by sites found in internal introns, for example in the *KCT13* gene, and these were also upregulated after ZFC3H1-depletion (Supplemental Figure 1C). When the levels of these IPAs were compared between the nucleus and the cytoplasm, we found that their cytoplasmic levels generally grew more than their nuclear levels, suggesting that ZFC3H1 was also required for their nuclear retention (Figure 1F). This could be clearly seen in individual examples (Figure 1G-I).

As documented previously, we also saw that ZFC3H1-depletion caused an upregulation in certain PROMPTs (an example is shown in Supplemental Figure 1D) and in many cases these tended to accumulate more rapidly in the cytoplasm than in the nucleus (an example is shown in Supplemental Figure 1E).

From these results we conclude that depletion of ZFC3H1 results in the upregulation of IPA transcripts and their accumulation in the cytoplasm. These results are inline with previously published results examining MTR4-depleted cells (Ogami et al. 2017). In addition, ZFC3H1-depletion led to the upregulation of a subset of PROMPTs, as described by others (Meola et al. 2016; Silla et al. 2020; Wu et al. 2020), and their cytoplasmic accumulation.

### Depletion of ZFC3H1 leads to the inhibition of nuclear export of mRNAs generated from distal 3’ polyadenylation/cleavage sites

Although ZFC3H1-depletion did not grossly affect the nuclear/cytoplasmic distribution of most mRNAs, we did notice that for mRNAs generated from genes with several 3’ polyadenylation/cleavage signals (Supplemental Figure 2A), the cytoplasmic/nuclear distribution of the long mRNA isoforms, generated from distal 3’ polyadenylation/cleavage sites, tended to be more nuclear in the ZFC3H1-depleted cells when compared to control cells (Supplemental Figure 2B-C). In contrast the overall distribution of mRNAs was generally unaffected (see “mRNAs” in Supplemental Figure 2B). To determine whether this inhibition of export was true for short isoforms produced from the same genes, we compared the portions of the 3’UTR that are common to both short and long isoforms (cUTR, see Supplemental Figure 2A). Reads that mapped to cUTR regions were less affected but still more skewed towards the nucleoplasmic fraction (Supplemental Figure 2B, D). Since these reads come from both long and short isoforms, we attempted to infer the short UTR counts by subtracting out the signal in the cUTR reads that came from the long isoform (see methods). The distribution of these inferred short UTR counts between the nucleus and cytoplasm were relatively unaffected (Supplemental Figure 2E). To determine whether these trends were due to an overall destabilization of long isoforms we next compared the change in cytoplasmic/nuclear distribution to the mRNA’s overall levels. We observed that upon ZFC3H1-depletion the decrease cytoplasmic/nuclear distribution was much larger than any change in levels for the long isoform mRNAs (Supplemental Figure 2F). Again, this trend was less pronounced in cUTR reads (which come from both short and long isoforms), and completely disappeared in the inferred short UTR counts (Supplemental Figures 2G-H). These trends were also seen when the levels in the cytoplasm and nucleus were directly compared with total levels (Supplemental Figure 3).

From these results we conclude that ZFC3H1 is likely required for the export of mRNAs generated from distal 3’ polyadenylation/cleavage sites.

### ZFC3H1, but not other members of the PAXT complex, are required for the nuclear retention of reporter mRNAs containing 5’SS motifs

Previously, we documented that mRNAs that contain intact 5’SS motifs were nuclear retained. Since these generally resemble IPA transcripts, we next tested whether MTR4 or ZFC3H1 were required for their nuclear retention. We thus treated U2OS cells with lentiviral delivered shRNAs against MTR4 or ZFC3H1, or control shRNAs (Figure 2A) and transfected plasmids containing the reporter mRNA *fushi tarazu* (*ftz*) with and without the 5’SS motif (Figure 2B) as previously described (Lee et al. 2015, 2020). Note that the 5’SS motif is present in a DNA region found in many expression vectors and encodes a V5-His epitope tag; however, within the *ftz* reporter this region is present downstream of the stop codon and is not translated. Additionally, this version of *ftz* lacks any intron (*Δi*). To successfully deplete MTR4 we used a pool of two separate shRNAs, as transduction with any single lentiviral-delivered shRNA gave poor depletion (data not shown). In contrast to what had been seen previously in HeLa cells (Ogami et al. 2017; Silla et al. 2018), MTR4-depletion did not lead to the co-depletion of ZFC3H1 (Figure 2A). This allowed us to determine whether MTR4 directly affects IPAs independent of ZFC3H1. Depletion of ZFC3H1 was performed using two independent shRNAs, and this did not affect MTR4 levels (Figure 2A).

When we analyzed the distribution of the *ftz* reporter mRNA by fluorescent *in situ* hybridization (FISH), we found that MTR4-depletion did not affect the cytoplasmic/nuclear levels of *ftz* mRNA containing the 5’SS motif (*ftz-Δi-V5-His*) (Figure 2C-D). In contrast, depletion of ZFC3H1 with either shRNA led to a significant inhibition in the nuclear retention of the same mRNA (Figure 3C-D). Depletion of either MTR4 or ZFC3H1 had no effect on the cytoplasmic/nuclear levels of *ftz* lacking the 5’SS motif (*ftz-Δi*) (Figure 3C-D).

**Figure 3.**
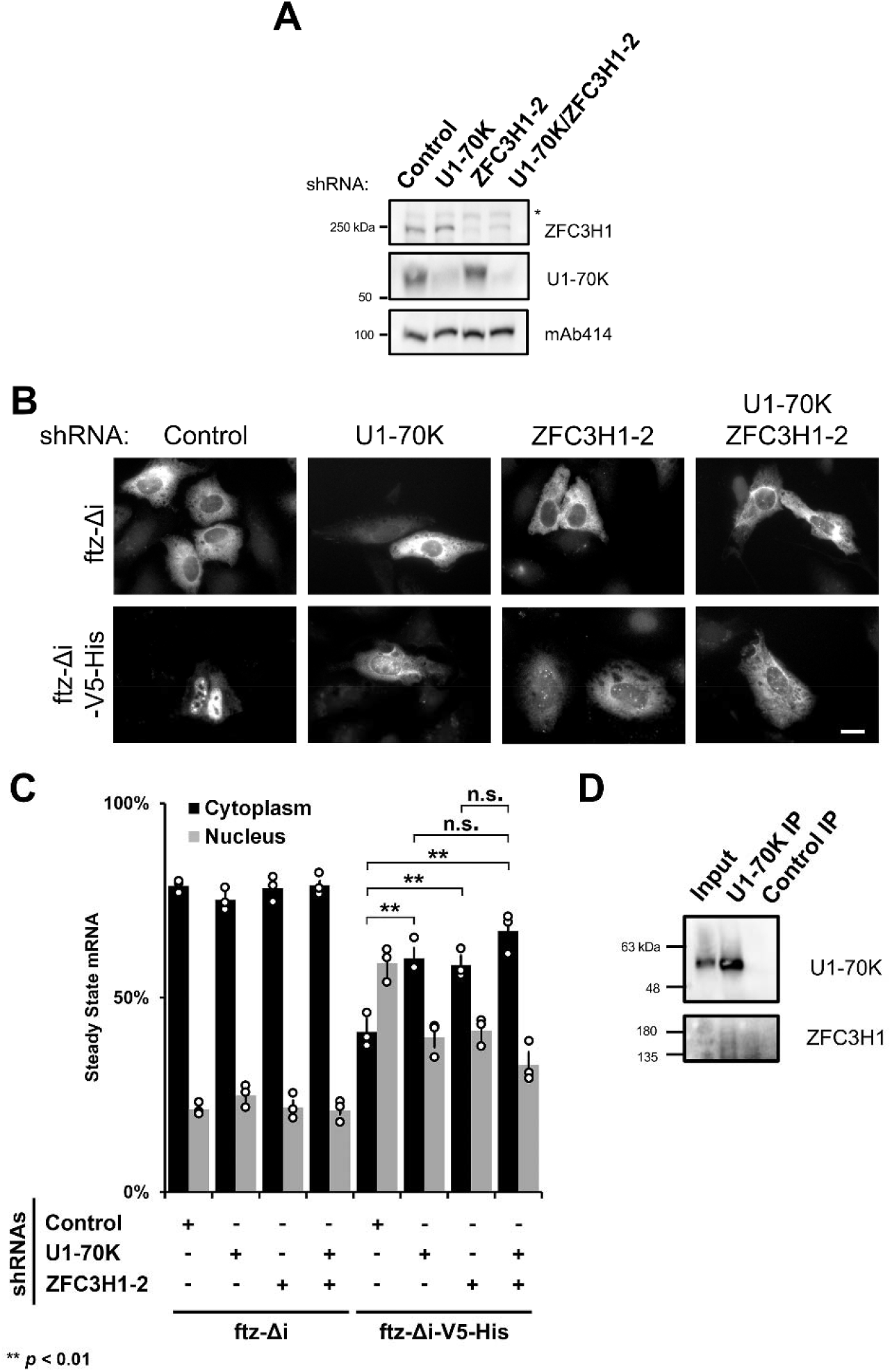
ZFC3H1 and U1-70K function in the same pathway for the nuclear retention of 5’SS motif containing mRNAs. (A) U2OS cells were treated with lentivirus shRNA against either U1-70K, ZFC3H1 or a mixture of the two. Lysates were collected 96 hrs, separated by SDS-PAGE and immunoprobed for U1-70K, ZFC3H1 and mAb414. Note that to effectively deplete u1-70K, cells were treated with lentivirus containing four shRNA plasmids. Also note that the asterisk (*) denotes a non-specific band. (B-C) Control-, U1-70K-, ZFC3H1- or co-depleted cells were transfected with the intronless *ftz* reporter +/- *V5-His* and as for figure 2. The cytoplasmic localization of *ftz-*Δ*i-V5-His* mRNA in cells co-depleted of U1-70K and ZFC3H1 resembles the single depletion, suggesting that both proteins function in the same pathway. Representative images are shown in (C), scale bar = 10 µm, and quantification is shown in (D). Each bar represents the average and standard error of at least three independent experiments, each experiment consisting of at least 30 to 60 cells. Student t-test was performed for (C), * *p* < 0.05, ** *p* < 0.01, *** *p* < 0.001. (D) U2OS cell lyates were subjected to immunoprecipitation reactions with mouse anti-U1-70K antibody (“U1-70K IP”), or mouse IgG (“Control IP”). Immunoprecipitates were separated by SDS-PAGE and immunoprobed for U1-70K and ZFC3H1. For comparison, a small amount of input lysates (3%) was also analyzed. Note that the predominant ZFC3H1 band appears to migrate faster than in samples that are rapidly denatured (see A), likely due to proteolytic cleavage after extended incubations.

Next, we analyzed the effect of PABPN1-depletion on the distribution of our reporter mRNAs. We depleted PABPN1 with two independent shRNAs in U2OS cells (Supplemental Figure 4A) and monitored the distribution of various *ftz* mRNA reporters (Supplemental Figure 4B) by FISH. We did not see a significant change in the cytoplasmic/nuclear distributions of *ftz* mRNA containing the 5’SS motif. This was true for intronless (*ftz-Δi-V5-His*) or intron-containing (*ftz-i-V5-His*) versions of *ftz* (Supplemental Figure 4C-D). We did see a modest, but significant inhibition in the nuclear export of our control reporter (*ftz-Δi*), in line with previous findings that PABPN1 is required for efficient mRNA nuclear export (Apponi et al. 2010). Thus, it remains formerly possible that since overall mRNA nuclear export is lower in PABPN1-depleted cells, this could have counteracted any putative increase in cytoplasmic levels of the 5’SS motif-containing mRNAs due to a general inhibition of nuclear retention. Previously, we showed that a general inhibition in mRNA export by depleting the nuclear pore protein TPR led to a further enhancement in the nuclear retention of *ftz-Δi-V5-His* mRNA (Lee et al. 2020), unlike what we see after PABPN1-depletion. These considerations not withstanding, our results suggest that PABPN1-depletion does not have a large impact on the nuclear retention of reporter mRNAs containing 5’SS motifs.

### mRNA nuclear retention is not dependent on alterations in 3’ cleavage or poly(A)-tail length

Interestingly, ZFC3H1 was identified in a mass spectrometry analysis of U1 snRNP-associated factors, which also contained 3’ processing components (So et al. 2019). Additionally, the presence of 5’SS motifs has been shown to inhibit both 3’ cleavage and polyadenylation (Boelens et al. 1993; Gunderson et al. 1998; Vagner et al. 2000). In light of these observations, *ftz-Δi-V5-His* mRNA might be nuclear retained due to alterations in 3’ cleavage or 3’ processing that are potentially regulated by ZFC3H1.

To determine whether the presence of a 5’SS motif disrupted normal 3’ cleavage or polyadenylation of the *ftz* mRNA reporter, we expressed intron lacking and containing versions of this reporter, with and without the 5’SS motif in U2OS cells, extracted mRNA from these lysates and assessed the 3’ cleavage by 3’ RACE and the poly(A)-tail length using the ePAT assay (Figure 2E) (Janicke et al. 2012). By 3’ RACE, both *ftz-Δi* and *ftz-Δi-V5-His* had the same 3’ cleavage site (Figure 2F) indicating that the 5’SS did not alter the 3’ end of the transcript. We also found that the presence of the V5-His region did not alter the size of the ePAT amplicons (Figure 2G), which comprises the 3’ end of the RNA and the entire poly(A)-tail, which we estimate to be ∼120 nucleotides long (Figure 2E). Although, it is likely that in many cases the presence of a 5’SS affects 3’ cleavage, this may depend on the strength of the 3’ cleavage/polyadenylation signal. In cases where these signals are strong, as in the *ftz* reporter which contains the bovine growth hormone polyadenylation signal (Pfarr et al. 1986), cleavage and polyadenylation are likely not affected. Thus, we conclude that the ability of the 5’SS to inhibit nuclear export is functionally distinct from its ability to inhibit 3’ cleavage and polyadenylation. This finding is in agreement with the observation that the PAXT complex acts on PROMPTs after they have become polyadenylated (Wu et al. 2020).

### U1-70K acts in the same pathway as ZFC3H1 in the nuclear retention of 5’SS motif containing mRNAs

As described above, ZFC3H1 was identified in a mass spectrometry analysis of U1 snRNP-interacting proteins (So et al. 2019). In addition, immunoprecipitates of ZFC3H1 contained U1-70K, a component of the U1 snRNP as detected by mass spectrometry (Meola et al. 2016). These observations raised the possibility that 5’SS motifs are initially recognized by U1 snRNP and that in the absence of a 3’SS motif, which would normally recruit other components of the spliceosome, U1 recruited ZFC3H1 to redirect the transcript to a nuclear retention and decay pathway. To assess whether U1 snRNP was required for the nuclear retention of 5’SS motif containing mRNAs, we used a mixture of four lentiviral delivered shRNA to deplete U1-70K (Figure 3A). U1-70K has been shown to be required for many activities of the U1 snRNP outside of splicing, including polyadenylation and 3’ cleavage (Gunderson et al. 1998; Vagner et al. 2000). We observed that U1-70K-depleted U2OS cells did not efficiently retain *ftz* reporters that contained a 5’SS motif (*ftz-Δi-V5-His*) in the nucleus (Figure 3B-C). In contrast, U1-70K-depletion had no effect on the distribution of *ftz-Δi* mRNA, which lacks a 5’SS (Figure 3B-C). Note that ZFC3H1 was unaffected by U1-70K-depletion.

To test whether these factors functioned in the same pathway, we assessed the distribution of our reporter mRNAs in cells depleted of both ZFC3H1 and U1-70K and compared these to the single depletions (Figure 3A). We observed that the double depleted cells had cytoplasmic/nuclear levels of *ftz-Δi-V5-His* that were not significantly different than in each of the single depleted cells (Figure 3B-C). Note that the distribution of this reporter was still more nuclear than a version that lacked a 5’SS motif (compare *ftz-Δi-V5-His* to *ftz-Δi* Figure 3C). These data strongly indicate that nuclear retention of *ftz-Δi-V5-His* was only partially inhibited in the single and double knockdown cells and that ZFC3H1 and U1-70K act in the same pathway.

We next sought to confirm previous mass spectrometry reports that suggested an interaction between U1 and ZFC3H1 (Meola et al. 2016; So et al. 2019). Indeed, we detected endogenous ZFC3H1 in immunoprecipitates of endogenous U1-70K but not in control immunoprecipitates (Figure 3D). These results indicate that ZFC3H1 and U1-70K are both in the same complex in cells.

In summary, these data suggest that both ZFC3H1 and U1-70K (and by extension the U1 snRNP) retain 5’SS motif containing mRNAs using a common pathway, and likely by forming a complex.

### ZFC3H1 is required for the egress of 5’SS motif containing mRNAs from nuclear speckles

Previously, we showed that 5’SS motif containing mRNAs that are retained in the nucleus also co-localize with nuclear speckles and remain in these compartments under steady state conditions (Lee et al. 2015). To determine if ZFC3H1 or U1-70K are required for targeting newly synthesized mRNAs with 5’SS motifs to nuclear speckles, we microinjected plasmids containing either the intronless β-globin gene without (β*G-Δi*) or with (β*G-Δi-V5-His*) a 5’SS motif into the nuclei of cells depleted of ZFC3H1 or U1-70K, or treated with control shRNA. We then fixed cells at various time points and monitored the co-localization of newly synthesized mRNA with the nuclear speckle marker SC35 by Pearson correlation coefficient analysis, as we have done previously (Akef et al. 2013; Lee et al. 2015). Note that microinjection allows for the generation of a large amount of mRNA in a short time span allowing one to observe events that happen only in the early stages of gene expression (Gueroussov et al. 2010). Since the *ftz* mRNA is trafficked to speckles independently of the 5’SS (Akef et al. 2013), we used the β*G-Δi* mRNA which requires the 5’SS motif for nuclear speckle localization (Supplemental Figure 5A), as previously described (Akef et al. 2013; Lee et al. 2015).

In control shRNA-treated cells, we observed that within 30 min of expression, β*G-Δi-V5-His* mRNA began to accumulate into nuclear speckles and the degree of speckle localization increased over time (Supplemental Figure 5B-C). In contrast, β*G-Δi* mRNA was not as robustly targeted to speckles during this time course. The speckle targeting β*G-Δi-V5-His* mRNA was not affected by depletion of ZFC3H1 or U1-70K (Supplemental Figure 5B-C). We also monitored the speckle targeting of *ftz-Δi* without and with a 5’SS motif. Again, depletion of ZFC3H1 did not significantly impact the targeting of this mRNA to nuclear speckles (Supplemental Figure 5D).

We next investigated whether ZFC3H1 was required for the egress of mRNAs containing 5’SS motifs out of nuclear speckles. To determine this, we monitored the localization of *ftz* without (*ftz-Δi*) and with (*ftz-Δi-V5-His*) a 5’SS motif in transfected cells where the total amount of nuclear speckle localization reflects steady state dynamics (i.e. both nuclear speckle targeting and egress). Note that we did not use β*G-Δi-V5-His* mRNA for these experiments, as the β*G-Δi* mRNA contains other elements that inhibit nuclear export and promote decay in a nuclear speckle-independent manner (Akef et al. 2015). We found that depletion of ZFC3H1 resulted in a decrease in nuclear speckle localized mRNA by Pearson correlation coefficient analysis (Figure 4A-B). When we quantified the amount of mRNA in nuclear speckles, the 5’SS motif caused an increase, and this was reversed in cells depleted of ZFC3H1 (Figure 4C).

**Figure 4.**
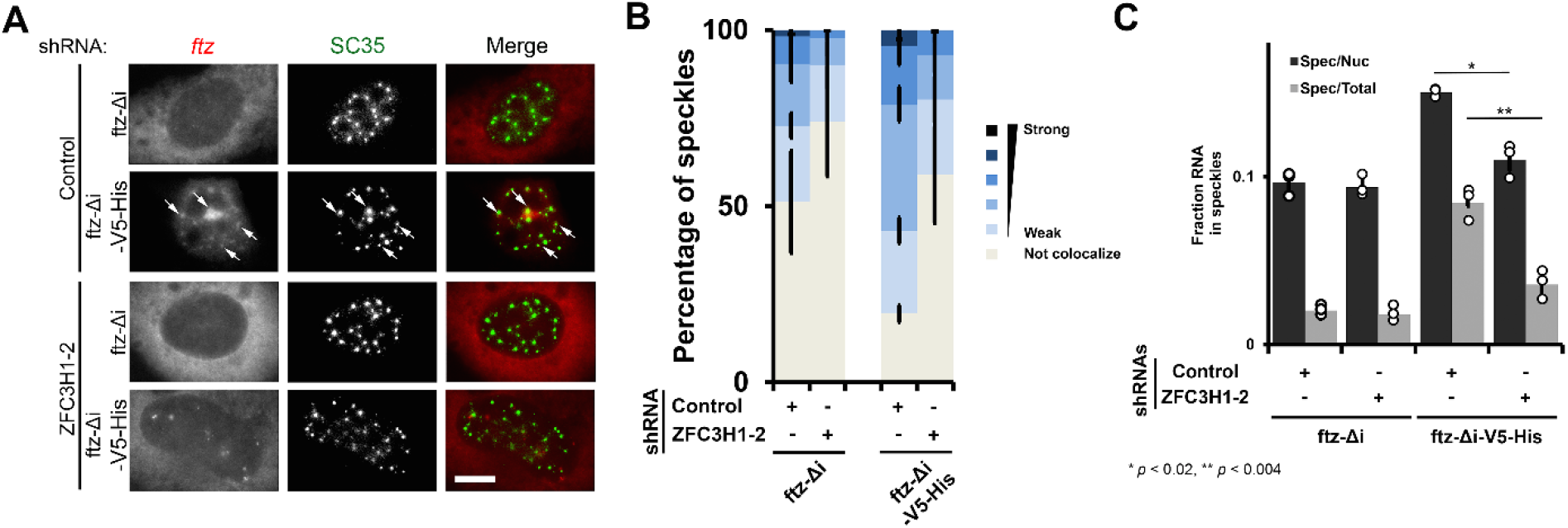
ZFC3H1 is required for nuclear retention of 5’SS motif containing mRNAs in speckles. (A-C) Control or ZFC3H1-depleted U2OS cells were transfected with *ftz-Δi +/- V5-His*. 18-24 hrs post-transfection, the cells were fixed and stained for *ftz* mRNA by FISH and for the nuclear speckle marker SC35 by immunofluorescence. Representative images, with each row depicting a single field of view, is shown in (A) with merged overlays showing *ftz* mRNA in red and SC35 in green. Scale bar = 10 µM. Examples of *ftz*/SC35 co-localization are indicated with arrows. (B) The degree of *ftz*/SC35 co-localization by Pearson correlation coefficient analysis was quantified as previously described (Akef et al. 2013), except that ‘not colocalized’ values < 0.25 and each bin contains values increasing by increments of 0.25. Note that ZFC3H1-depletion leads to decreased level of colocalization between *ftz* mRNA and SC35. Each bar represents the average and standard error of three independent experiments, each experiment consisting of 100-200 nuclear speckles from 10-20 cells. (C) The amount of *ftz* reporter mRNA present in nuclear speckles as a percentage of either the total nuclear (“spec/nuc”) or total cellular (“spec/total”) mRNA levels in transfected cells. Each data point represents the average and standard error of the mean of three independent experiments, each experiment consisting of 10-20 cells. Student’s t-test was performed, * *p* < 0.02, ** *p* < 0.004.

From these experiments we concluded that ZFC3H1 helps to promote the localization of 5’SS motif containing mRNAs to nuclear speckles.

### Nuclear speckles are required for the efficient nuclear retention of 5’SS motif containing mRNAs

Next we examined whether nuclear speckles are required for the nuclear retention of 5’SS motif containing mRNAs. We overexpressed GFP-CLK3 to trigger nuclear speckle disassembly, as was previously described (Wong et al. 2011), and examined the distribution of either *ftz-Δi-V5-His* and *ftz-Δi* mRNA, and of the speckle marker SC35. As a control we overexpressed Histone 1B-GFP (H1B-GFP). As was reported previously (Wong et al. 2011), expression of GFP-CLK3 resulted in the disappearance of detectable nuclear speckles (Figure 5A-C), and in these cells we observed that *ftz-Δi-V5-His* mRNA was significantly enriched in the cytoplasm (Figure 5A, D). In some cells the nuclear speckle marker, SC35 completely disappeared, while in other cells its levels did not significantly drop, but it became diffuse throughout the nucleoplasm. In H1B-GFP expressing cells, *ftz-Δi-V5-His* mRNA was nuclear, similar to what we had seen in control U2OS cells (Figure 5A, C). Cells expressing GFP-CLK3 still exported *ftz-Δi*, which lacks the 5’SS motif, indicating that expression of this kinase did not disrupt normal mRNA export (Figure 5C-D, Supplemental Figure 6A).

**Figure 5.**
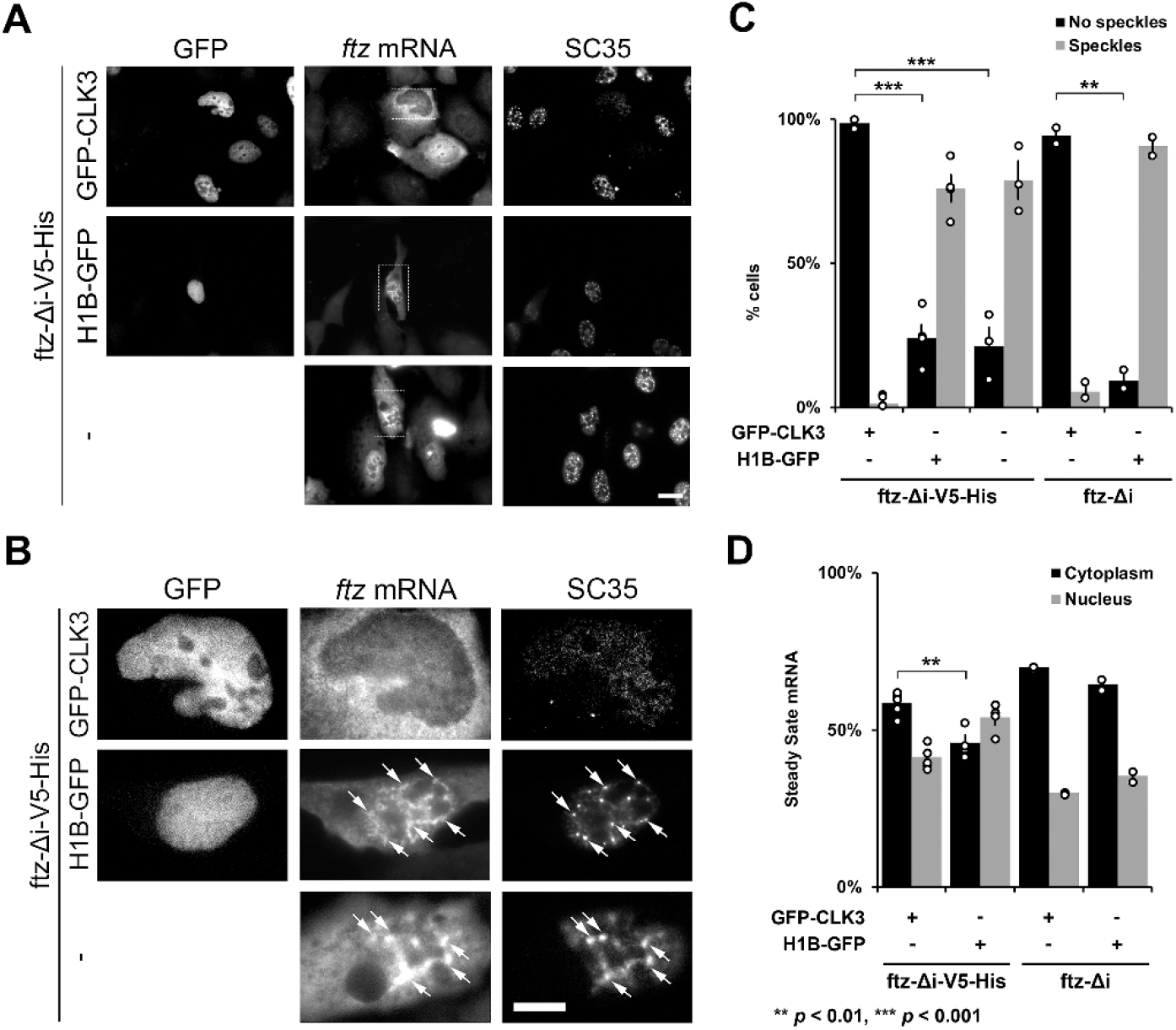
Nuclear speckles promote mRNA nuclear retention of 5’SS motif containing mRNA. (A-D) U2OS cells were transfected with *ftz-Δi-V5-His* or *ftz-Δi*, alone (“-“) or with either GFP-CLK3, or H1B-GFP. 24 hrs post-transfection, cells were fixed and stained for *ftz* mRNA by FISH and for the speckle marker SC35 by immunofluorescence. Representative images, with each row depicting a single field of view imaged for GFP, *ftz* mRNA and SC35 is shown in (A). Note that the overexpression of GFP-CLK3 disrupts nuclear speckles (SC35) compared to H1B-GFP or non GFP-expressing cells. Scale bar = 10 µM. Enlarged images of the regions indicated in (A) are depicted in (B). Examples of *ftz*/SC35 co-localization are indicated with arrows. Scale bar = 10 µM. (C) The number of cells that lacked or contained nuclear speckles were quantified. Each bar represents the average and standard error of two (*ftz-Δi*) or four (*ftz-Δi-V5-His*) independent experiments, each consisting of 30-50 cells. Note that expression of GFP-CLK3 disrupts nuclear speckles compared to control cells. *** *p* < 0.001 (Student’s t-test). (D) Quantification of the cytoplasmic/nuclear FISH signal with each bar representing the average and standard error of two (*ftz-Δi*) or four (*ftz-Δi-V5-His*) independent experiments, each experiment consisting of at least 30 to 60 cells. ** *p* > 0.01 (Student’s t-test).

Interestingly, expression of *ftz-Δi-V5-His* alone caused many cells to have larger nuclear speckles than what is typically seen in untransfected cells (Figure 5B, Supplemental Figure 6B), likely because these regions grew as more and more mRNA became incorporated into them. Co-expression of H1B-GFP did cause some cells to have smaller nuclear speckles than what is seen in untransfected cells, however most cells still contained these structures (Figure 5B, also see Supplemental Figure 6B).

In summary, our results suggest that nuclear speckles are required for the efficient nuclear retention of 5’SS motif containing mRNAs, although it remains possible that GFP-CLK3 expression disrupts mRNA nuclear retention independently of its effects on nuclear speckles.

## Discussion

Here we show that 5’SS motif containing mRNAs require ZFC3H1 and U1-70K for their efficient nuclear retention. Moreover, our data indicates that these two proteins act in the same pathway. It is likely that when a newly synthesized transcript has a 5’SS motif, it first recruits the U1 snRNP to initiate splicing, however, if this is followed by a strong 3’cleavage/polyadenylation signal ZFC3H1 is recruited instead. This leads to nuclear retention in nuclear speckles and degradation of the transcript. In support of this we demonstrate that upon ZFC3H1-depletion, there is a widespread stabilization and export of endogenous IPA transcripts, which contain intact 5’SS motifs followed by poly(A)-tails.

Interestingly, depletion of MTR4 and PABPN1 had minimal effects on the nuclear retention of reporter mRNAs containing 5’SS motifs in U2OS cells. This is in contrast to previous findings in HeLa cells (Ogami et al. 2017). It remains possible that PAPBN1 disrupted both nuclear retention and export, complicating the interpretation of our results. Indeed, when mRNA export is disrupted by TPR-depletion, we observed that *ftz-Δi-V5-His* mRNA was even more nuclear (Lee et al. 2020), unlike our current results in PAPBN1-depleted cells. As for the discrepancy between our results with MTR4 and past results, this is likely explained by the fact that MTR4-depletion caused co-depletion of ZFC3H1 in HeLa (Ogami et al. 2017; Silla et al. 2018) but not U2OS cells (Figure 2A). It is possible that other factors present act redundantly with MTR4 in U2OS cells. Alternatively, nuclear retention of IPA transcripts may only require low levels of MTR4.

There has been much interest in the role of nuclear speckles in promoting nuclear retention and export (Akef et al. 2013; Dias et al. 2010; Jain and Vale 2017; Palazzo and Lee 2018; Wang et al. 2018). Our new results show that ZFC3H1 reduces the nuclear speckle egress of mRNAs. Interestingly, this was true for both mRNAs with and without 5’SS motifs (Figure 4B). We still do not understand how these mRNAs are targeted to speckles. Likely this is due in part to U1 snRNP, although not U1-70K. It remains possible that U1 snRNP, which remains intact after U1-70K-depletion (Rösel-Hillgärtner et al. 2013), can still target mRNAs to speckles but fails to recruit ZFC3H1 to the transcript. This is in agreement with previous results where the tethering of U1-70K to a reporter mRNA promoted nuclear retention without targeting the transcript to nuclear speckles (Takemura et al. 2011).

Intriguingly, in the fission yeast *S. pombe*, homologs of ZFC3H1 (Red1) and MTR4 (Mtl1) are required for both the decay and the sequestration of meiotic mRNAs and transposable element derived transcripts into nuclear foci (Shichino et al. 2018, 2020; Sugiyama and Sugioka-Sugiyama 2011). These results suggest that the ability of ZFC3H1 to negatively regulate expression of mRNAs and ncRNAs, by promoting retention in nuclear subdomains and triggering RNA decay, is widely conserved in eukaryotes.

Previously, we showed that mRNAs with 5’SS motifs still acquired TREX components, such as UAP56, and the nuclear transport receptor NXF1 but fail to be exported (Lee et al. 2015). Thus, it is likely that the export-promoting activity of these factors is inhibited by ZFC3H1. Indeed, it appears that UAP56-dependent ATP-hydrolysis is required for nuclear egress (Hondele et al. 2019) and this is directly counteracted by ZFC3H1.

In previously published results, we found that microinjected mRNAs with 5’SS motifs were efficiently exported from nuclei (Lee et al. 2015). This may indicate that U1 snRNP and/or ZFC3H1 may be recruited during transcription, and this is supported by observations that U1 snRNP directly associates with RNA polymerase II (Zhang et al. 2021). However, since microinjected mRNAs are spliced within 15 mins after being injected into nuclei (Palazzo et al. 2007), it is likely that microinjected mRNAs are still recognized by U1 snRNP but not ZFC3H1. This suggests that ZFC3H1 may be recruited to nascent transcripts by the joint action of U1 and RNA polymerase II in a co-transcriptional manner.

Our RNA Frac-Seq data indicates that ZFC3H1 likely inhibits the nuclear export of PROMPTs and other ncRNAs, as indicated by previous reports (Silla et al. 2018), and also promotes the export of mRNAs generated from distal 3’cleavage/polyadenylation sites (Supplemental Figures 2 and 3). It may be that this later class of mRNAs are outcompeted by ncRNAs and IPAs for binding to nuclear export factors. Alternatively, ZFC3H1 may directly participate in their nuclear export. Further studies will be needed to test these various possibilities.

## Material and methods

### Plasmids constructs and antibodies

All plasmids for reporter constructs in pcDNA3.0 and H1B-GFP were previously described (Akef et al. 2013; Lee et al. 2015, 2020; Palazzo et al. 2007). Plasmid expressing GFP-CLK3 was previously described (Wong et al. 2011).

Antibodies used in this study include rabbit polyclonals against ZFC3H1 (also known as CCDC131) (Bethyl laboratories, A301-457A), MTR4 (also known as SKIV2L2) (Bethyl laboratories, A300-614A), PABPN1 (Bethyl laboratories, A303-523A), U1-70K (Abcam, ab83306), Aly (Zhou et al. 2000) and TRAPα (Görlich et al. 1990) or mouse monoclonals against U1-70K (Sigma-Aldrich, clone 9C4.1), mAb414 (Sigma), SC35 (Clone SC35, Sigma), and α-tubulin (DM1A, Sigma). Immunofluorescence with Alexa647-conjgated secondaries (Thermo-Fisher) and HRP-conjugated secondaries (Cell Signaling). All antibodies were diluted 1:1000 for western blotting and 1:100 to 1:250 for immunofluorescence microscopy.

### Cell culture, DNA transfection experiments and Lentiviral delivered shRNA protein depletion

U2OS and HEK293T cells were grown in DMEM media (Wisent) supplemented with 10% fetal bovine serum (FBS) (Wisent) and 5% penicillin/streptomycin (Wisent). DNA transfection experiments were performed as previously described (Akef et al. 2013; Lee et al. 2015, 2020; Palazzo et al. 2007).

For all DNA transfections, U2OS cells were transfected with the appropriate amount of DNA plasmid according to the manufacturer’s protocol using GenJet U2OS DNA *in vitro* transfection reagent (SignaGen Laboratories) for 18 to 24 hrs.

The lentiviral delivered shRNA protein depletion was performed as previously described (Akef et al. 2013; Lee et al. 2015, 2020). Briefly, HEK293T were plated at 50% confluency on 60 mm dishes and transiently transfected with the gene specific shRNA pLKO.1 plasmid (Sigma), packaging plasmid (*Δ* 8.9) and envelope (VSVG) vectors using Lipo293T DNA in vitro transfection reagent (SignaGen Laboratories) according to the manufacture’s protocol. 48 h post-transfection, viruses were harvested from the media and added to U2OS cells pre-treated with 8 μg/ml hexadimethrine bromide. Cells were selected with 2 μg/ml puromycin media for at least 4 to 6 days. Western blotting was used to determine the efficiency of ZFC3H1, MTR4, PABPN1 and U1-70K depletion. The shRNAs constructs (Sigma) used in this study are as follows: ZFC3H1-1 “TRCN0000129932” 5’-CCGGGCCAAG AAGCAATCTA TCAATCTCGA GATTGATAGA TTGCTTCTTG GCTTTTTTG-3’, ZFC3H1-2 “TRCN0000432333” 5’-CCGGGACTGA TGACATCGCT AATTTCTCGA GAAATTAGCG ATGTCATCAG TCTTTTTTG-3’, MTR4-1 “TRCN0000307086” 5’-CCGGCCCAGG ATAGAAGAGT CAATACTCGA GTATTGACTC TTCTATCCTG GGTTTTTG-3’, MTR4-2 “TRCN0000296268” 5’-CCGGAGCAGG ACCACTTCGT CAAATCTCGA GATTTGACGA AGTGGTCCTG CTTTTTTG-3’, PABPN1-1 “TRCN0000000124” 5’-CCGGAGGTAG AGAAGCAGAT GAATACTCGA GTATTCATCT GCTTCTCTAC CTTTTTT-3’, PABPN1-2 “TRCN0000000120” 5’-CCGGCCCATA ACTAACTGCT GAGGACTCGA GTCCTCAGCA GTTAGTTATG GGTTTTT-3’, U1-70K-A “TRCN0000000011” 5’-CCGGCCAAGG GTAGGTGTCT CATTTCTCGA GAAATGAGAC ACCTACCCTT GGTTTTT-3’, U1-70K-B “TRCN0000349622” 5’-CCGGGACATG CACTCCGCTT ACAAACTCGA GTTTGTAAGC GGAGTGCATG TCTTTTTG-3’, U1-70K-C “TRCN0000287201” 5’-CCGGGCACCA TACATCCGAG AGTTTCTCGA GAAACTCTCG GATGTATGGT GCTTTTTG-3’ and U1-70K-D “TRCN0000287138” 5’-CCGGCGATGC CTTCAAGACT CTGTTCTCGA GAACAGAGTC TTGAAGGCAT CGTTTTTG-3’.

### Microinjections, fluorescent in situ hybridization (FISH) staining, immunostaining and nuclear speckle Pearson correlation and enrichment quantifications

Microinjections, fluorescent in situ hybridization and immunostaining were performed as previously described (Gueroussov et al. 2010; Lee and Palazzo 2017; Lee et al. 2020).

Nuclear speckle localization by Pearson correlation was performed as previously described (Akef et al. 2013; Lee et al. 2015, 2020). Briefly, for each cell the 10 brightest nuclear speckles, as observed by SC35 staining were selected and Pearson correlation analysis was conducted between the FISH signal and SC35 immunofluorescence signal using areas rectangular regions of interest 1□4 μm^2^ in size. Each experiment consists of an analysis of 10 cells (for a total of 100 speckles). Graphs in Figure 4 and Supplemental Figure 5 consist of the average and standard deviation of 3 independent experiments.

The quantification of mRNA enrichment in speckles was performed as previously described (Akef et al. 2013). Briefly, thresholds were drawn on the SC35 immunofluorescence channel so that 10% (+/− 0.5%) of the nuclear area was selected per cell. Using this selected area, the fluorescence intensity of RNA was calculated and divided by either the total integrated mRNA signals in the nucleus (“Spec/Nuc”) or the cell body (“Spec/Total”).

### 3’RACE, RT-PCR and ePAT

For 3’RACE and RT-PCR experiments, total RNA was extracted from transfected U2OS cells using Trizol (Life Sciences). About 1 μg total RNA was used for first strand synthesized using murine MLV reverse transcriptase (Invitrogen) and oligo(dT) primer according to the manufacturer’s instructions. For 3’RACE experiments, the resulting cDNA was amplified using *ftz* F’ primer 5’-ATG GGG TGT TGT CCC GGC TGT TGT - 3’ and an oligo(dT) primer. The resulting PCR product was inserted into CloneJet (Fermentas) vector following the sticky-end cloning protocol (see manufacture’s instructions) and transformed into DH5α competent cells. DNA was extracted from colonies using Miniprep kit (Geneaid) and sent for sequencing.

ePAT was performed as previously described (Janicke et al. 2012). Total RNA was extracted by Trizol from cell transfected with the indicated *ftz* plasmid and expressed for 18-24 hrs. The ePAT anchor primer used was 5’-GCG AGC TCC GCG GCC GCG TTT TTT TTT TTT-3’, the universal primer used was 5’-GCG AGC TCC GCG GCC GCG-3’ and *ftz*-specific ePAT primer used was 5’-ATG GGG TGT TGT CCC GGC TGT TGT - 3’.

### Co-IP experiment

∼ 1.2 10^7^ U2OS cells were trypsinized, pelleted and washed 3X times in ice-cold 1X PBS and flash frozen in liquid nitrogen. The cell pellet was lysed in ∼1.4 ml of IP buffer (20 mM Tris-HCl, pH8, 137 mM NaCl, 1% NP-40, 2 mM EDTA and *cOmplete* mini-protease inhibitor (Roche)) mixed for 25 minutes at 4 °C to ensure complete lysis. To ensure that the nuclear proteins were released from the chromatin, the sample was passed through a 25¾” needle to sheer the chromatin multiple times before and after incubation. The cell lysis was cleared by centrifugation at 16,100*g* for 10 min and 0.65 ml supernatant was mixed with 50μl Protein G beads (NEB, #37478S) pre-conjugated to U1-70K antibody (Sigma-Aldrich, clone 9C4.1), pre-washed 3X times with IP wash buffer (same as IP buffer, except that 0.05 % NP-40 was used). The cell lysate was incubated for 3 hours at 4 °C. As a control experiment, the same amount of cell lysate was added to the same volume of or Mouse IgG antibody (Santa Cruz, sc-2025). Following incubation, the beads were washes 4 to 5X times with IP wash buffer. To elute the proteins, 50μl of 2.5X Laemmli sample buffer was added to all the samples and boiled for 5 minutes. Samples were separated on an SDS-PAGE gel and transferred onto a blot for immunoblotting.

### RNA Frac-Seq, data processing and analysis

RNA Frac-Seq were performed as previously described (Lee et al. 2020). RNA Fraq-Seq data were deposited in GEO database as GSE176144. IPA analysis was performed as described (Wang et al. 2019) using a curated list of IPA transcripts (Supplemental Table 1).

To estimate reads from short 3’UTRs, we used these calculations:

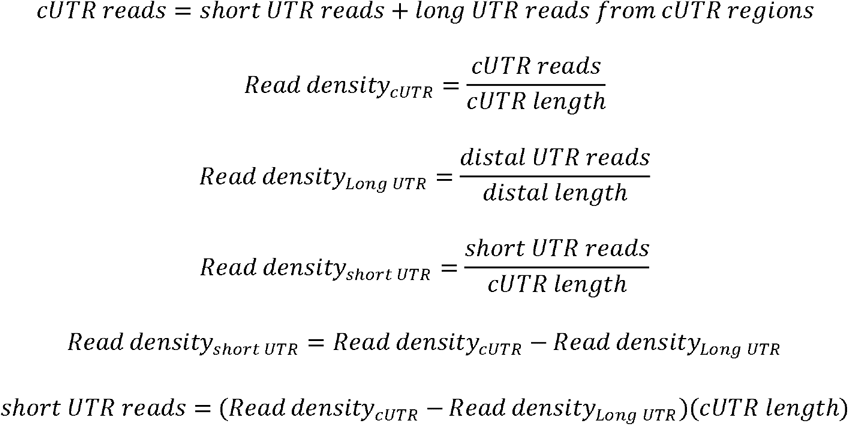

## Research Contribution

E.S.L. and H.W.S. performed the experiments under the guidance of A.F.P.

E.S.L. and A.F.P. co-wrote the paper. E.J.W. and A.G. performed the bioinformatics analysis under the guidance of A.F.P. and B. T.

## Funding

This work was supported by a grant from the Canadian Institutes of Health Research to AFP (FRN 102725).

## Conflict of Interest

The authors have declared that no conflict of interest exists.

## Acknowledgements

We thank T. Dubric and her staff at the Donnelly Centre Sequencing Facility for performing the RNA sequencing and quality control, and S. Ihn for providing critical feedback on the manuscript. We thank A. Cochran for generously providing us with GFP-CLK3 plasmid.

## Supplementary Figures and Tables

**Figure S1.**
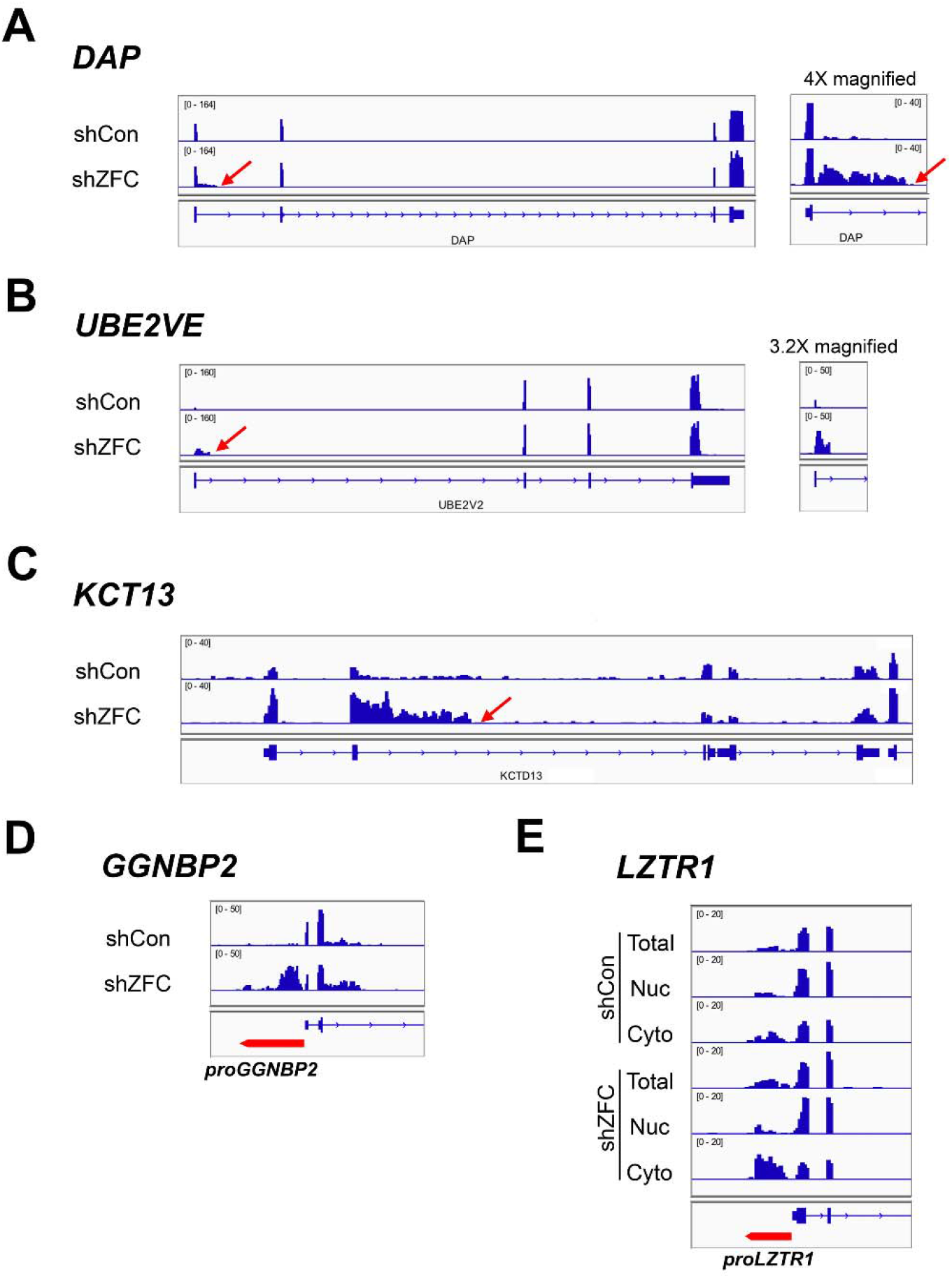
Genome browser tracks of endogenous 5’SS motif containing transcripts. (A-C) Selected examples of IPA transcripts upregulated following ZFC3H1-depletion in total RNA samples. The intronic 3’cleavage/polyadenylation site is denoted by the red arrows. (D-E) Selected example of PROMPTs (“*proGGNBP2*” and “*proLZTR1*”) that are upregulated following ZFC3H1-depletion. The PROMPTs are denoted by a red arrow. Note that following ZFC3H1 depletion, “*proLZTR1*” is more cytoplasmic.

**Figure S2.**
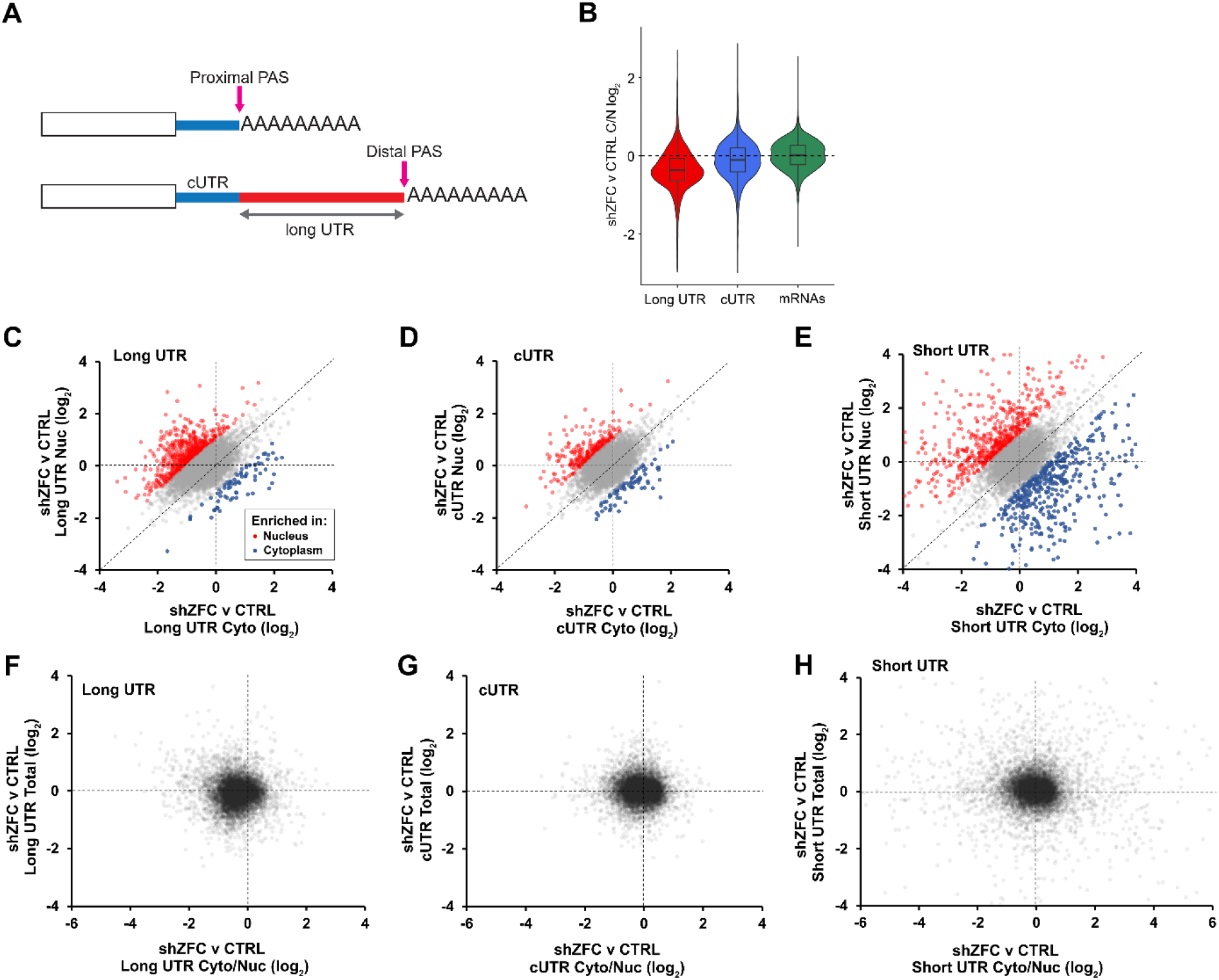
ZFC3H1-depletion leads to nuclear localization of mRNAs with long 3’ UTRs. (A) Schematic depicting alternative polyadenylation. If the proximal polyadenylation site (PAS) is used, this generates a short UTR isoform. If the distal PAS is used, this generates mRNAs with long 3’ UTRs (“long UTR”). Note that regions of the 3’UTR that are found only when the distal site is used are labeled in red while regions common to both short and long isoforms are denoted by “cUTR”. (B) Violin plots of the log_2_ change in cytoplasmic/nuclear ratio of mRNAs following ZFC3H1-depletion (“shZFC v CTRL C/N”) compared between reads mapping to long UTR regions (red regions in (A)), cUTR regions (blue regions in (A)) and all mRNA regions. Note that reads from long UTRs, which are generated from distal PASs, are more nuclear following ZFC3H1-depletion than reads mapping to cUTR or mRNAs in general. (C-E) The nuclear (*y-axis*)-cytoplasmic (*x-axis*) distribution of (C) long, (D) cUTR and (E) short UTRs following ZFC3H1-depletion. Each data point represents reads from a separate gene. See methods for how levels of short UTR reads were inferred. Note that ZFC3H1-depletion leads to nuclear retention of mRNAs with long UTRs (compare red dots to blue in (C)). This is not observed for short UTRs in (E). To eliminate the contribution of long UTRs from cUTR reads, we estimated reads that corresponds to only the short UTRs. (F-H) Similar to (C-E), except the cytoplasmic/nuclear ratio (*x-axis*) was compared to total RNA (*y-axis*). Note that the change in cytoplasmic/nuclear ratio for long UTR reads was much more pronounced than the change in total levels.

**Figure S3.**
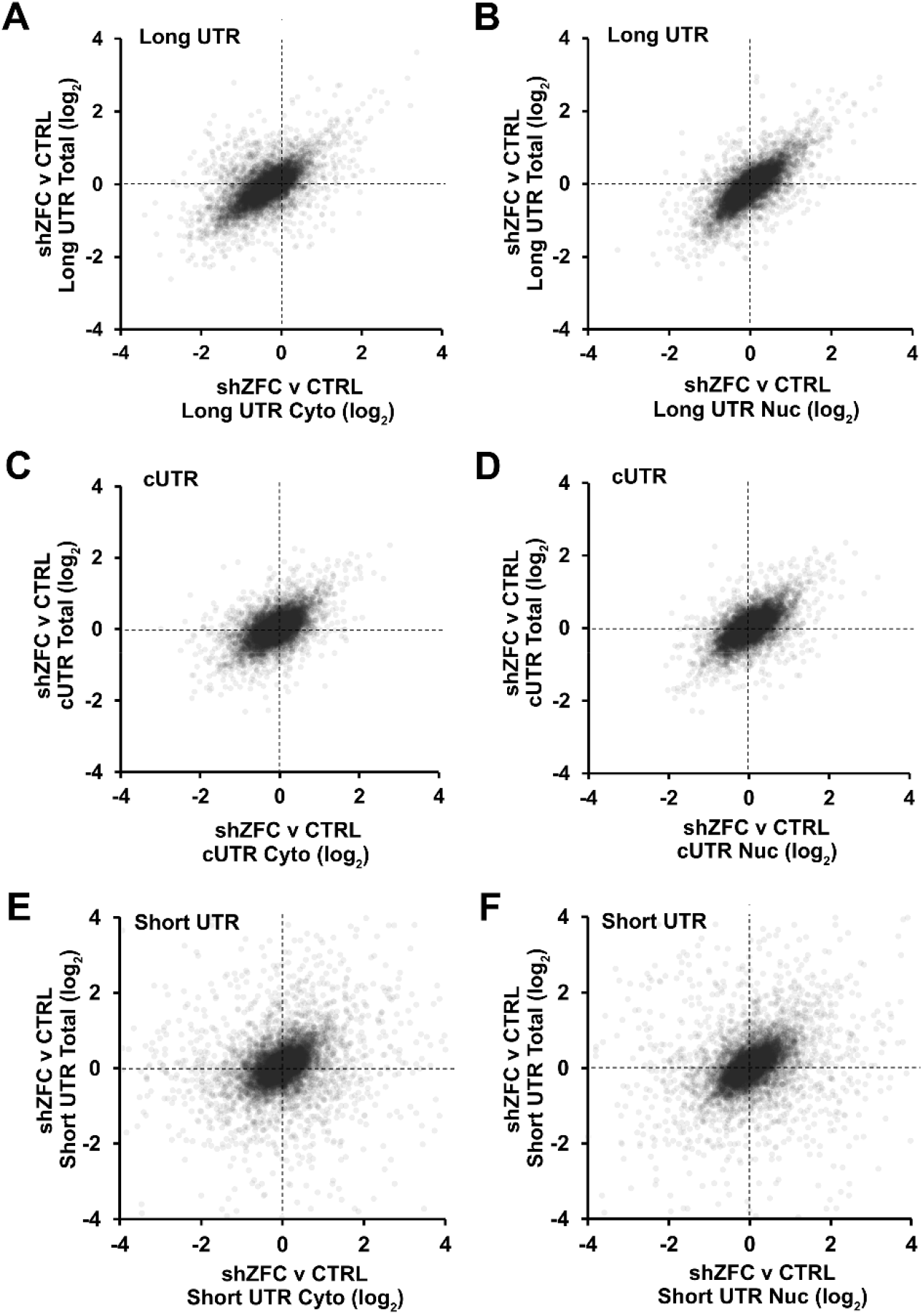
Analysis of nuclear and cytoplasmic levels of 3’UTR reads upon ZFC3H1-depletion. (A-B) Similar to Supplemental Figure 2C, except that the (A) cytoplasmic or (B) nuclear long UTR reads (*x-axis*) were plotted against total reads (*y-axis*) following ZFC3H1-depletion. (C-D) Similar to Supplemental Figure 2D, except that the (C) cytoplasmic or (D) nuclear long UTR reads (*x-axis*) were plotted against total reads (*y-axis*) following ZFC3H1-depletion. (E-F) Similar to Supplemental Figure S2E, except that the (E) cytoplasmic or (F) nuclear long UTR reads (*x-axis*) were plotted against total reads (*y-axis*) following ZFC3H1-depletion.

**Figure S4.**
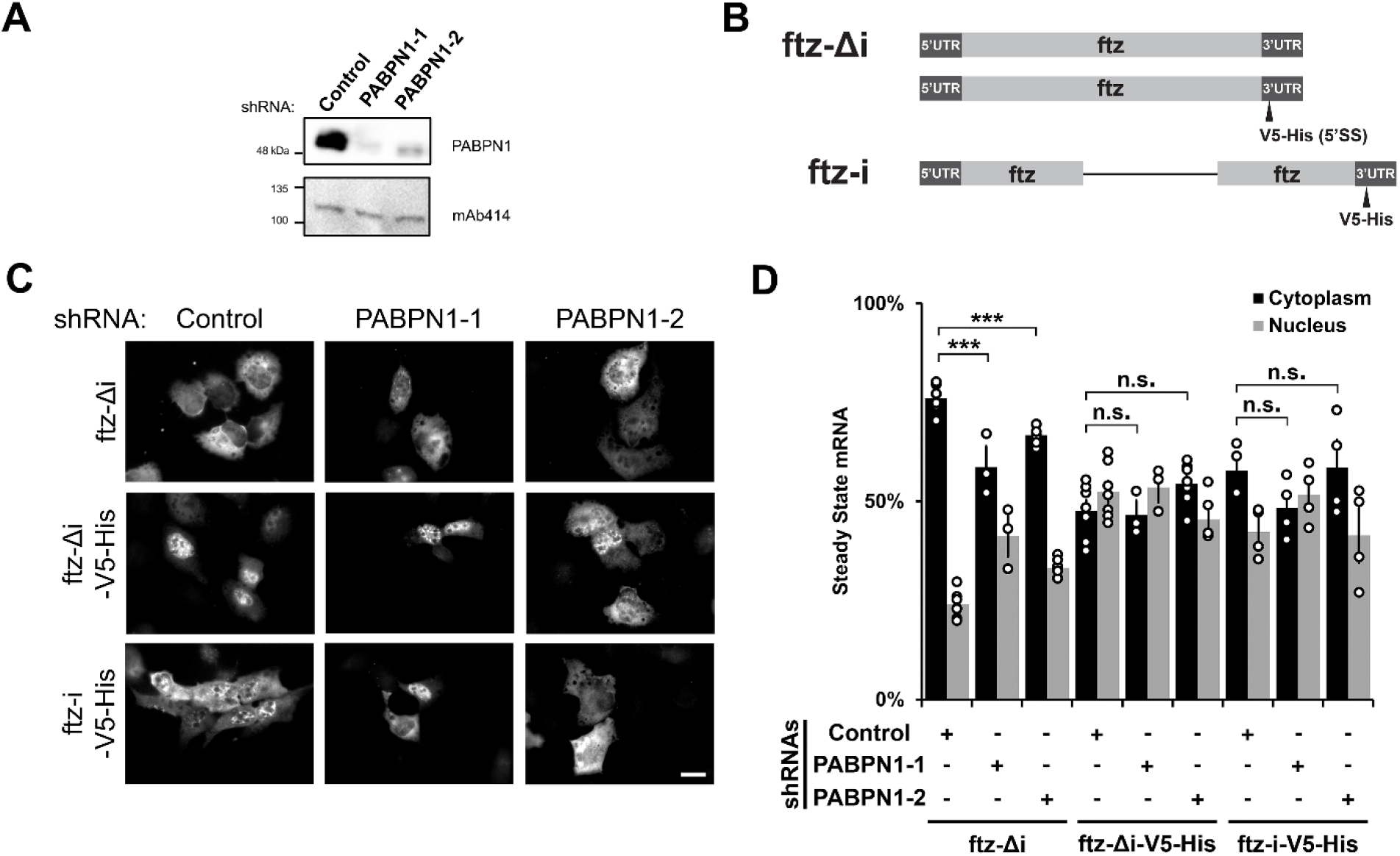
PABPN1-depletion has a minor effect on the nuclear retention of 5’SS motif containing mRNAs. (A) U2OS cells were treated with two different lentivirus shRNAs against PABPN1 (“PABPN1-1” and “PABPN1-2”). Lysates were collected after 72 to 96 hrs, separated by SDS-PAGE and immunoprobed for PABPN1 and mAb414, which recognizes FG-containing Nups (Davis and Blobel 1986). (B) Schematic of the *ftz* reporter constructs used. (C-D) Control- or PABPN1-depleted U2OS cells were transfected with the intronless *ftz* reporter +/- *V5-His* and the intron-containing *ftz-V5-His* reporter. After 18-24 hrs the cells were fixed, stained for *ftz*, imaged (C) and the cytoplasmic/nuclear ratios were quantified (D). Each bar represents the average and standard error of at least three independent experiments, each experiment consisting 30 to 60 cells. Student t-test was performed with *** denoting *p* < 0.001, N.S. not significant. Note that PABPN1-depletion reduced the cytoplasmic level of the control *ftz-Δi* mRNA but did not significantly affect the localization of the *ftz-V5-His* containing reporters. Scale bar = 10 µM.

**Figure S5.**
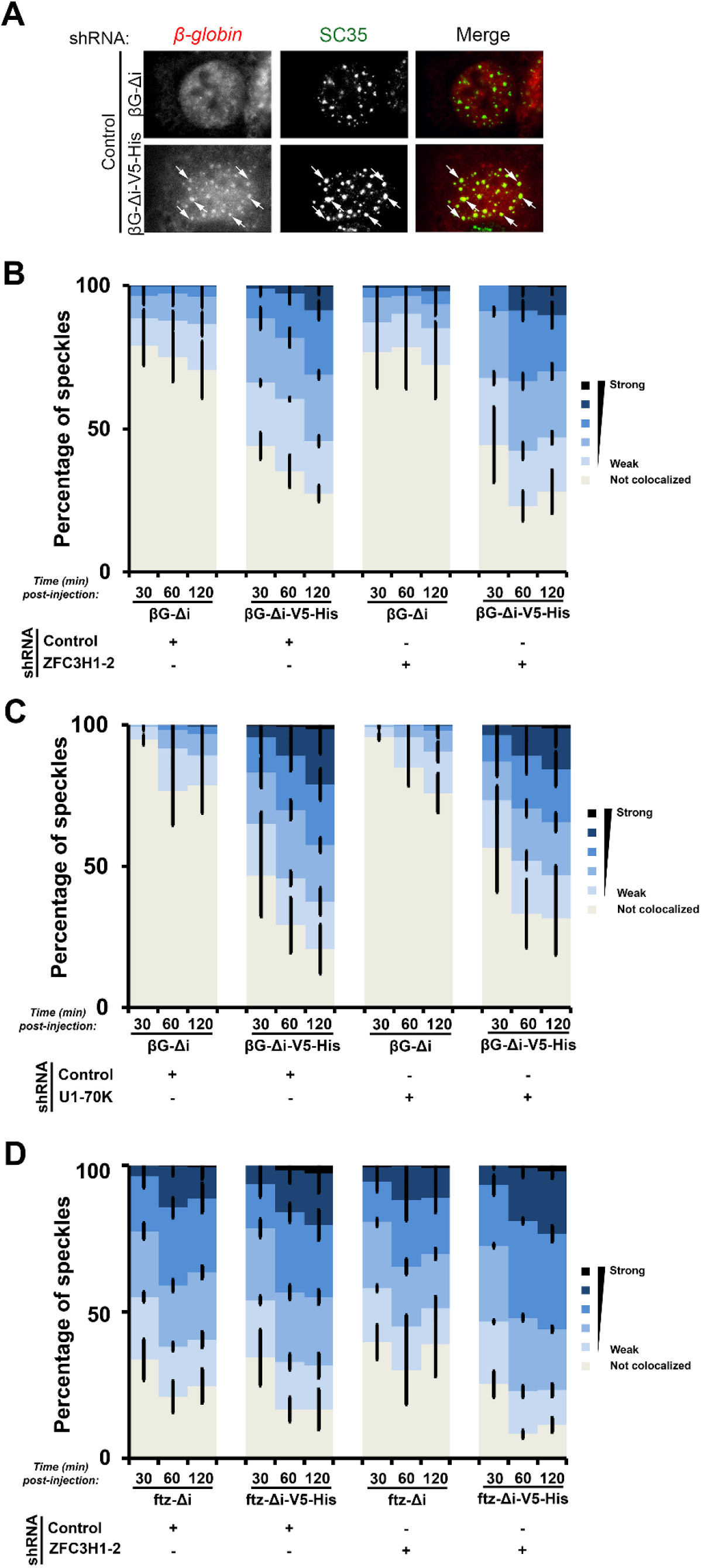
ZFC3H1 and U1-70K are not required for trafficking 5’SS motif containing mRNAs to nuclear speckles. Control, ZFC3H1- or U1-70K depleted U2OS cells were microinjected with plasmids containing the β*G-Δi +/- V5-His* (A-C) or *ftz-Δi +/- V5-His* (D) reporter plasmids. After the indicated times, the cells were fixed and stained for β*-globin* or *ftz* mRNA by FISH and for the nuclear speckle marker SC35 by immunofluorescence. (A) Example images of U2OS cells fixed 2 hours post-injection with β*-globin +/- V5-His* reporter mRNA, with each row representing a single field of view with white arrows point to examples of β*-globin*/SC35 co-localization. The merged overlayed image depicts β*-globin* mRNA in read, SC35 in green. Scale bar = 10 µM. (B-D) Quantification of the degree of β*-globin*/SC35 (B, C) or *ftz*/SC35 (D) co-localization in cells depleted of ZFC3H1 (B, D), U1-70K (C) or after control shRNA-treatment (B-D) by Pearson correlation coefficient analysis. Each bar represents the average and standard error of three independent experiments, each experiment consisting of 150 to 200 nuclear speckles from 15-20 cells. Note that depletion of either ZFC3H1 or U1-70K did not affect the targeting of 5’SS motif containing mRNAs to nuclear speckles.

**Figure S6.**
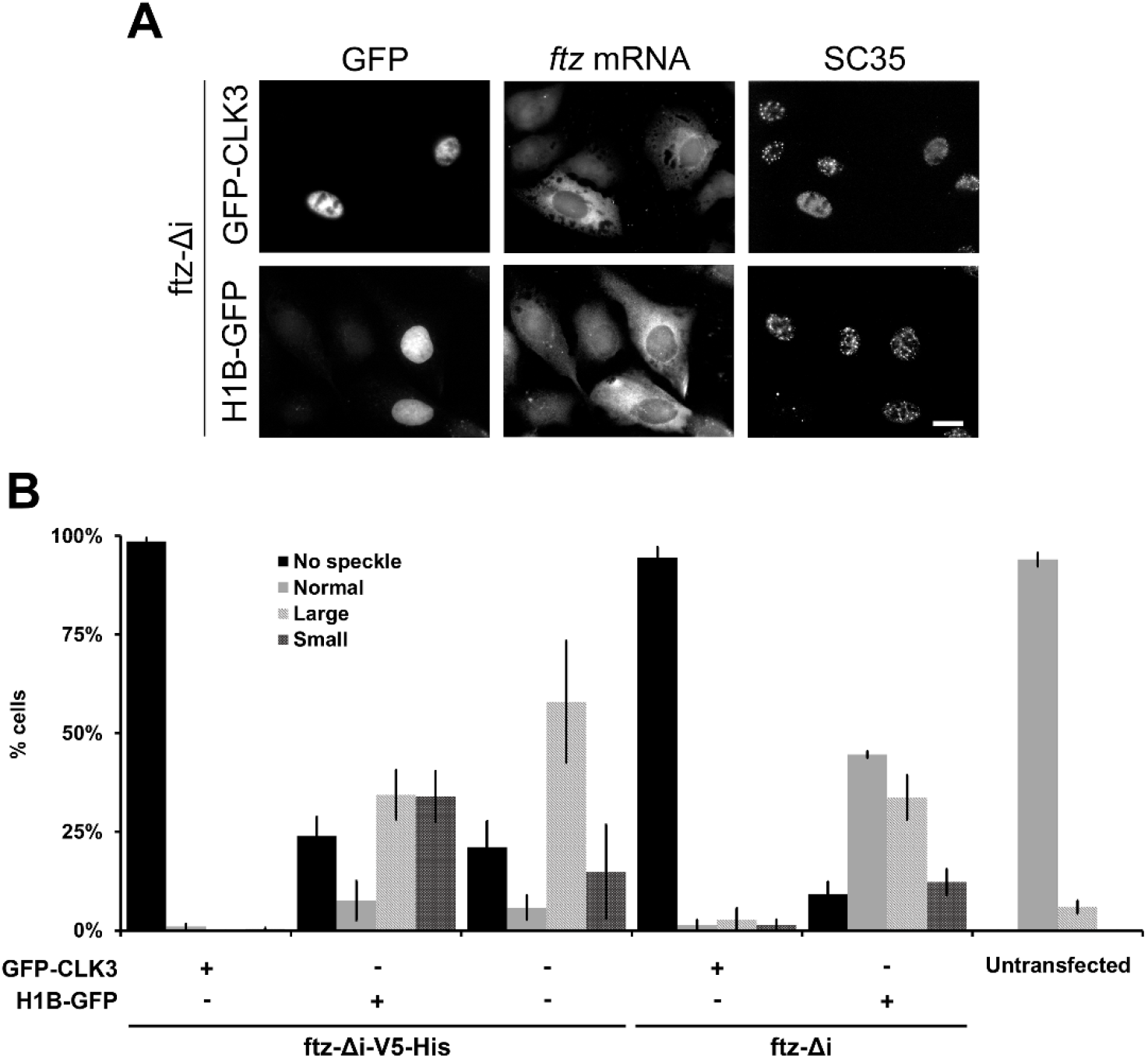
Disruption of nuclear speckles does not affect nuclear export of *ftz-Δi* mRNAs. (A) U2OS cells were co-transfected with *ftz-Δi* and GFP-CLK3 or H1B-GFP. 18 to 24 hrs post-transfection, cells were fixed and stained for *ftz* mRNA by FISH and for the speckle marker SC35 by immunofluorescence. Representative images, with each row depicting a single field of view imaged for GFP, *ftz* mRNA and SC35 is shown in (A). Scale bar = 10 µM. (B) U2OS cells were transfected with *ftz-Δi-V5-His* or *ftz-Δi* alone (“-“) or with either GFP-CLK3, or H1B-GFP. 24 hrs post-transfection, cells were fixed and stained for *ftz* mRNA by FISH and for the speckle marker SC35 by immunofluorescence and the nuclear speckle morphology was visually inspected and quantified for each cell. Untransfected cells were also quantified to determine the morphology of nuclear speckles without any exogenously expressed mRNAs of GFP-tagged proteins. Each bar represents the average and standard error of two (*ftz-Δi*) or four (*ftz-Δi-V5-His*) independent experiments, each consisting of 30-50 cells.

**Supplementary Table 1. List of IPA Transcripts Analyzed**.

## Notes

### Competing Interest Statement

The authors have declared no competing interest.

